## Supplementary figures and images for "Application of High Resolution Melt analysis (HRM) for screening haplotype variation in non-model plants: a case study of Honeybush (*Cyclopia* Vent.)"

### Supplementary Figure_Neighbor Joining clustering of wild C. subternata populations based on alleic diffreretiation (Jost's D)

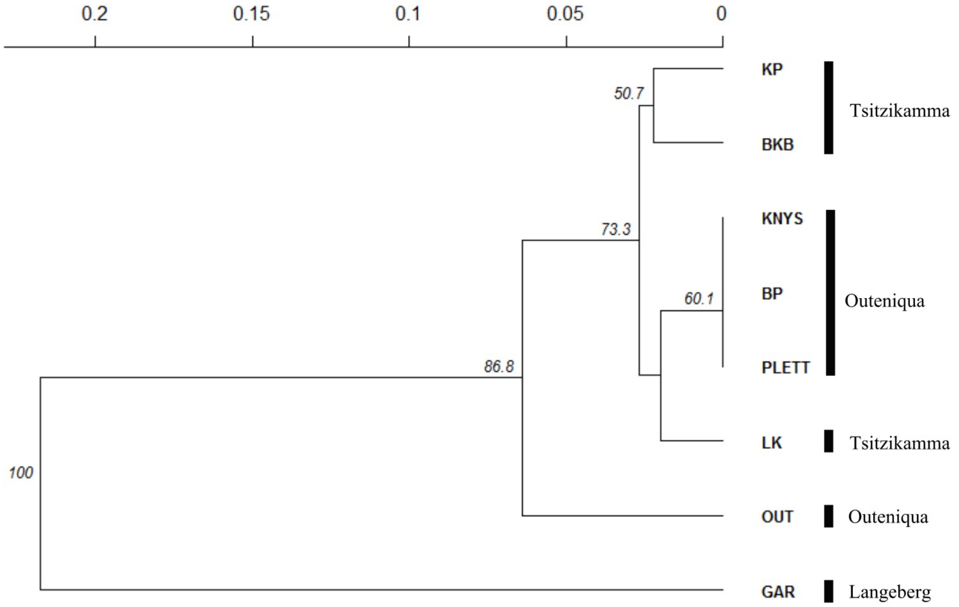

### Table S1_Species and regions initially screened for HRM primer design

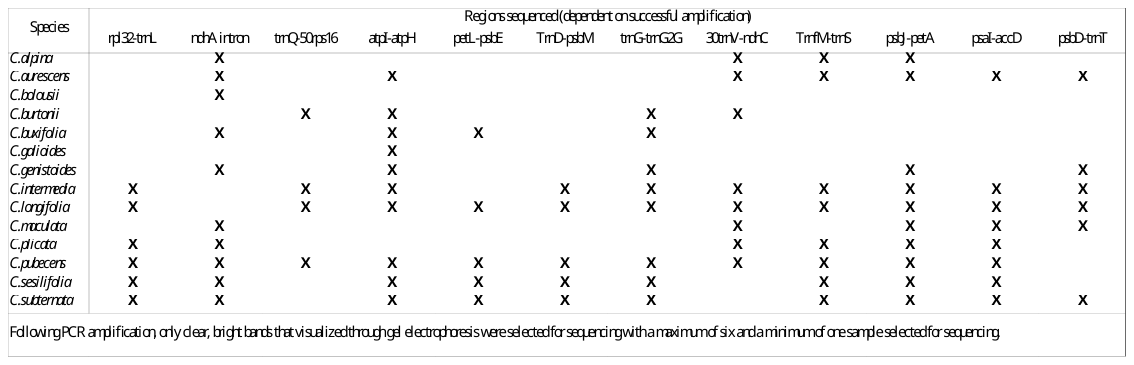

### Table S2_Genus-speific primers tested for amplification

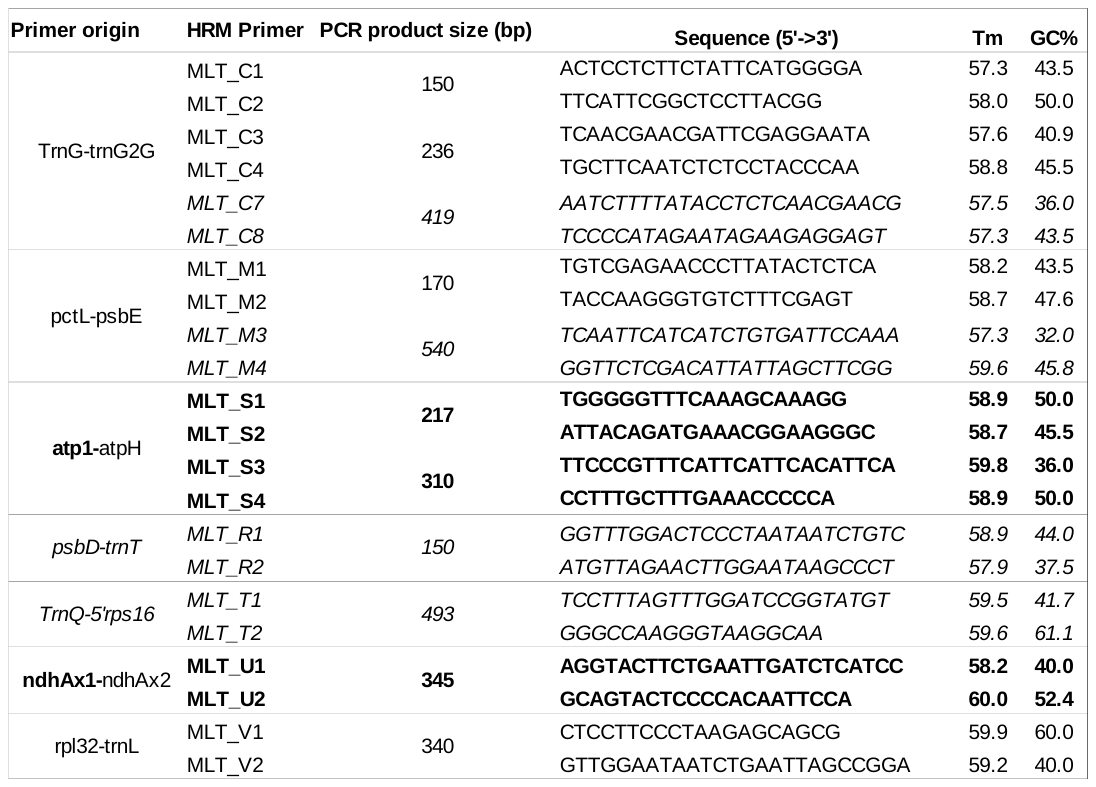

### Table S3_Nucleotide variation among haplotypes detected in wild C. subternata populations via HRM and confirmation Sanger sequencing

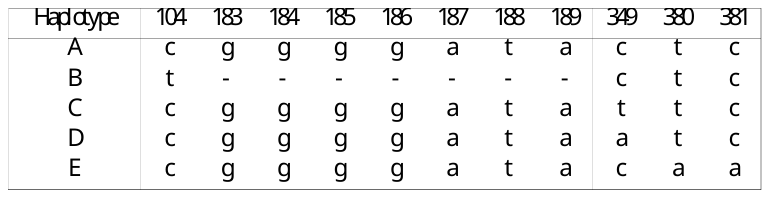

### Table S4_Costing analysis comparing HRM to alternative approaches to data generation

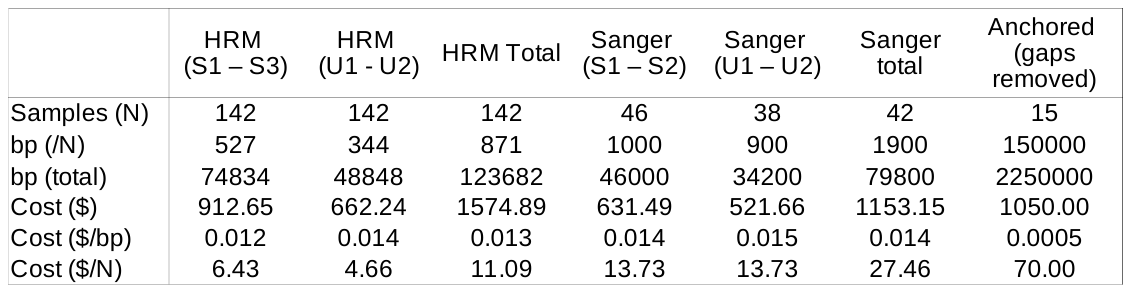
