## Supplementary Methods_Lab procedures for "Application of High Resolution Melt analysis (HRM) for screening haplotype variation in non-model plants: a case study of Honeybush (*Cyclopia* Vent.)"

### S1: lab protocols and R script

#### DNA extraction:

1: Grind plant material (+/- 0.02g). This is often done using liquid nitrogen in a pestle and mortar however, shaking dry plant material and glass beads in a 1.5ml flat bottomed screw cap reaction tube using a mixer-mill at highest frequency, much quicker.

2: Add 700ul preheated (65^O^C) PVP enriched 2x CTAB extraction buffer and 2ul of mercaptoethanol to each sample and vortex to ensure that all plant material is suspended in the buffer. The quantities of PVP and mercaptoethonol were doubled in order to remove the excess polyphenols present in *Cyclopia*.

3: Incubate the vortexed samples on a vibrating heating block at 65^O^C for an hour.

4: Add 600ul of chloroform:isoamyl alcohol (24:1, v/v) to each sample and mix by inversion for 5 min.

5: Centrifuge samples at 12 000 rpm for 5 min and pipette 600ul of the supernatant into a clean 1.5ul microfuge tube. Be sure to avoid pipetting any of the lower green liquid, rather collect less supernatant.

6: Add 5ul of RNase to each sample, mix by inversion and then incubate at 37^O^C for 15 min. (This step may be skipped, but is required for Next Generation sequencing).

7: Add 36ul of Ammoniumacetate and 600ul of ice cold ethanol:isopropanol (1:1, v/v) and mix by inversion before placing the samples into a freezer overnight to precipitate.

8: Centrifuge the chilled samples at 12 000 rpm for 5 minutes.

9: Pour out the ethanol:isopropanol and leave the tubes inverted for a short while (5 min) making sure not to loose the DNA pellet, use sterile paper towel to mop up any excess liquid.

10: Wash the DNA pellet in 250ul of 75% ethanol, generally the DNA pellet was dislodged and flatten out against the inside of the tube with the pipette tip before shaking the tube by hand. Be sure to replace the pipette tip between samples.

11: Centrifuge samples at 12 000 rpm for 5 min and then repeat steps 9 and 10 and centrifuge again.

12: Pour out the ethanol being careful not to loose your DNA pellet and then place the open tubes into an airtight container containing silica dessication crystals for 2-3 hours. Do not leave samples to dry in this manner overnight as it makes re-suspension difficult.

13: Re-suspend DNA in molecular grade water or TE buffer overnight and check the quality using your method of choice.

#### PCR primers from Shaw et al. 2007

Table A1: Polymerase Chain Reaction primers used to amplify the parent region of the loci screened by HRM. Primers in bold were used for unidirectional sequencing.

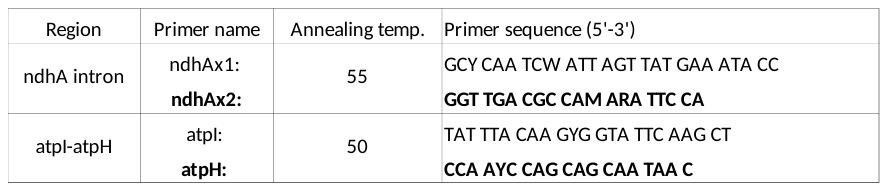

#### PCR protocol

Table A2: PCR amplification protocol for the primers sources from Shaw et al. 2007. See Table A1 for annealing temperatures. See
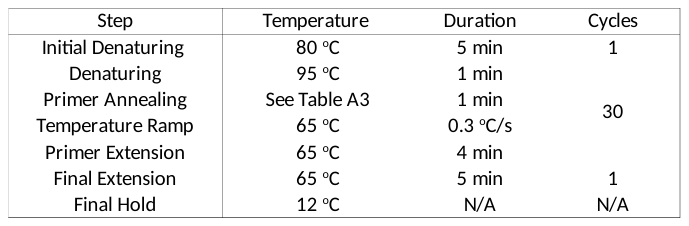

#### R script for assigning haplotypes to samples based on HRM clustering results

haps <-read.dna("Your haplotypes.fasta",format="fasta")

dat <- read.csv("Your sample haplotype classification.csv")

if(sum(!unique(dat$Haplotype)%in%rownames(haps))>0) warning("Some haplotypes in the .csv are NOT PRESENT IN THE FASTA FILE")

\# Removing haplotypes in the csv not present in the fasta file

dat <- subset(dat,Haplotype%in%rownames(haps))

for (i in 1:nrow(dat)) {

if (i==1) {

newalignment <- haps[dat[i,"Haplotype"],]

} else {

newalignment <- rbind(newalignment,haps[dat[i,"Haplotype"],])

}

}

rownames(newalignment) <- apply(dat,1,paste,collapse="_")

#check the alignment of the output in condon!!!
